# Comparative transcriptome revealed potential genes related with drug resistance in Melanoma

**DOI:** 10.1101/2022.12.14.520500

**Authors:** Bei Zhao, Yinghua Liu, Xuemei Tang, Shi Cheng

## Abstract

Chemotherapy remains a relatively ineffective and unsatisfactory treatment because of drug resistance in melanoma (whether due to intrinsic resistance or the use of cytostatic drugs). In order to explore the genes and signaling pathways related to melanoma drug resistance, the study presented here obtained drug-resistant melanoma cell lines of A375 and M14 by gradually increasing the concentration of dacarbazine (DTIC), followed by comparative transcriptomics studies (RNA-seq) and real-time quantitative PCR (RT-qPCR) and Western Blotting validation. The results showed that after 8 months of continuous treatment, the IC50 values of A375 and M14 to DTIC were increased by more than 5 folds. Meanwhile, flow cytometry analysis found that drug-resistant melanoma cells have a significant ability to resist apoptosis induced by DTIC. Subsequently, RNA-seq revealed high expression of *SENP1* and abnormal activation of the Hippo signaling pathway in drug-resistant cells. Finally, we found that compared with wild-type cells, the expressions of *SENP1* and *YAP* were both up-regulated in drug-resistant cells via RT-qPCR and Western Blotting. Roles of SENP1 in drug resistance was finally verified via its overexpression in normal A375 cell lines. Therefore, this paper infers that there is a positive correlation between the ubiquitin-specific protease SENP1 and the drug resistance of melanoma. Meanwhile, the up-regulation of its expression may lead to changes in the Hippo signaling pathway and increase the resistance of melanoma to DTIC.

## Introduction

Melanoma is a skin cancer caused by malignant lesions of melanocytes and has become the fastest growing cancer in men and the second fastest growing cancer in women (Kalal et al., 2017). According to statistics, there were 324,635 new cases of skin melanoma worldwide in 2020, of which 57,043 finally lost their lives (Sung et al., 2021). In fact, most melanomas are curable if diagnosed early. Nevertheless, the prognosis is poor once melanoma metastasizes, even accounting for more than 80% of the deaths caused by skin cancer (Wu and Singh, 2011). The median survival time of advanced patients is only 7.5 months, along with a rate of about 15% for the 2-year survival and only 5% for the 5-year survival (Mattila et al., 2018).

For early-stage skin melanoma, surgery is the main treatment method. Additionally, for patients with middle-advanced melanoma, adjuvant therapy such as chemotherapy or palliative therapy is an important means to improve the prognosis. Among them, dacarbazine (DTIC) is a conventional chemotherapy drug approved by the US FDA for the treatment of advanced melanoma (Velho, 2012). Even so, chemotherapy is currently a relatively ineffective and unsatisfactory treatment because of drug resistance in melanoma (whether due to intrinsic resistance or the use of cytostatic drugs), making it a major problem limiting chemotherapy. Therefore, studying the drug resistance mechanism of melanoma becomes the key to solving the problem.

In order to further explore the genes and signaling pathways related to melanoma drug resistance, the study presented here took A375 and M14 melanoma cells as the research object, obtaining drug-resistant cells by gradually increasing the concentration of DTIC added. The parent and drug-resistant melanoma cell line were then analyzed through comparative transcriptomics to explore the significantly differentially expressed genes and signaling pathways. Finally, the overexpression of related gene proved its association with drug resistance in melanoma. Our study will provide new insights about the drug resistance mechanisms of melanoma.

## Materials and Methods

### Cell lines and reagents

In this experiment, A375 and M14 cell lines used were preserved by the laboratory of Sichuan Provincial People’s Hospital, China. The main reagents used in this study include: dacarbazine (DTIC, D129847, Aladdin, China), Annexin V-EGFP/PI Cell Apoptosis Detection Kit (TransGen, Beijing, China), dimethyl sulfoxide (DMSO, Solarbio, Beijing, China), skimmed milk powder (Solarbio, Beijing, China), TBST buffer (Solarbio, Beijing, China), phosphate buffer (Solarbio, Beijing, China), RIPA lysate (Solarbio, Beijing, China), N,N′-methylenebisacrylamide (Aladdin, Shanghai, China), tetramethylethylenediamine (TEMED, Aladdin, Shanghai, China), ammonium persulfate (Aladdin, Shanghai, China), Coomassie Brilliant Blue (Aladdin, Shanghai, China), SYBR qPCR Master Mix (Vazyme, Nanjing, China).

### Instruments

The instruments used in this study included Multifunctional microplate detector (Synergy HTX, Biotek, USA), gel imaging system (ChemiDoc™ XRS+, Bio-rad, USA), flow cytometer (FACSCalibur, BD Biosciences, USA) and real-time quantitative PCR (QuantStudio ™ 3 Real-Time PCR System, ABI, USA).

### Apoptosis detection

Cell apoptosis was detected by flow cytometry. The cells in the control group and the experimental group were inoculated in 24-well plates (5×10^4^ cells/well). After 48 hours of culture, the cells were collected (centrifuged at 1000 rpm for 5 minutes), washed once with phosphate buffer, and then added with 100 μl Annevix Binding Buffer. After that, 5 μl Annevix FITC and 5 μl PI were added successively and kept in dark at room temperature for 15 min. Finally, apoptosis detection was performed on flow cytometry after adding 150 μl Annevix Binding Buffer.

### Transcriptomic analysis

The wild-type and successfully established drug-resistant melanoma cells were rinsed with 4°C pre-cooled 1×PBS for 2 to 3 times, and then Trizol was added to isolate total RNAs. cDNA libraries were amplified and sequenced using the BGISEQ-500 platform (BGI Group, Shenzhen, China). FastQC (version 0.11.8) were used to check raw sequencing reads. Adapter trimming and low-quality filtering was analyzed by Cutadapt (v2.0) to get the clean data. The gene expression was quantified by featureCounts (v1.6.0), and differential expressed genes (DEGs) between treatment groups were created by DESeq2 (R version 3.3.2). Cutoff value of fold change ≥1.5 and adjust *P*-value ≤0.05 was used. The results were analyzed for gene expression levels between different groups, prediction of new transcripts, GO/KEGG enrichment analysis, protein interaction network analysis, and gene co-expression network construction.

### Real-time quantitative PCR (RT-qPCR) analysis

Real-time quantitative PCR was used to verify the mRNA level of the target gene in the control group and the experimental group, and to detect whether it was in line with the change in transcription level. 1 μL of each dilution was used as template for following qRT-PCR reaction. The qPCR reaction was carried out in 10 μL reactions containing 5 μL of qPCR Master Mix (Vazyme, Nanjing, China), 3 μL dH_2_O, 1 μL diluted template cDNA and 1 μL of each PCR primer, employing the QuantStudio ™ 3 Real-Time PCR System (Applied Biosystems, CA, USA) (Sun et al., 2017). Three technical replicates were performed for each condition. Data analysis was carried out using the QuantStudio 3 software (Applied Biosystems, CA, USA) and the 2^−ΔΔCT^ method (Livak and Schmittgen, 2001)

### Construction of gene-overexpressed cell lines

The third-generation Lentivirus lentivirus packaging system was utilized to transfect pCMV-R+pMDG+target gene to HEK293T cells. The culture medium was replaced after 12 hours of infection with A375 melanoma cells. The medium was replaced with fresh medium containing puromycin (200 ng/mL), and screen for 14 days to establish a stable expression cell line. The protein expression level of SENP1 was detected by WB.

### Western Blot (WB)

The effect of different treatment factors on the protein level was detected by WB. Cells with different treatment factors were collected, lysed with RIPA strong lysate, and total protein was extracted and quantified with an NanoDrop® ND-1000 UV-Vis Spectrophotometer. 50 μg of total protein taken from each sample was electrophoresed and transferred to a membrane [polyvinylidene fluoride Ethylene (polyvinylidene fluoride, PVDF) membrane], following by block with 5% skimmed milk powder/washing buffer (TBST buffer) on a shaker at room temperature for 1 h. The primary antibody was bound overnight at 4°C then washed 3 times with TBST, followed by secondary antibody binding and incubate at room temperature on a shaker for 1 h. After that, the sample was washed 3 times with TBST and added with ECL chemiluminescence reagent for 1.5-2 min. Finally, the result was observed with a gel imaging system.

### Statistical Analysis

Statistical software SPSS 18.0 was used for data analysis. Measurement data were expressed as ±s. Comparisons among multiple groups were performed by one-way analysis of variance. Significant differences between two groups were performed by *t*-test, and *P*<0.05 was considered statistically significant.

## Results

### Construction and characterization of drug-resistant melanoma cell lines

To construct drug-resistant melanoma cell lines, the A375 and M14 cells growing in the logarithmic phase were treated with DTIC for 24 hours with an initial induction dose of 2 μg/mL. Additional 2 μg/mL DTIC was added until the cells grew to a density of about 70%-80% of the final density. After continuously adding DTIC to A375 and M14 melanoma cell lines for 8 months, we tested the tolerance of the two experimental groups and the control groups. Detailly, the parent cell lines were used as the control group (i.e., A375-WT and M14-WT) while the finally obtained drug-resistant cell line was the experimental group (<u>D</u>TIC-<u>R</u>esistant, A375-DR and M14-DR). First, the half maximal inhibitory concentration (IC50) of each cell line to DTIC was measured. As shown in Fig. 1A, the resistance to DTIC was significantly increased for both A375-DR and M14-DR melanoma cell lines after 8 months of treatment. Among them, the IC50 of the A375-DR increased from 22.5±2.1 μM to 138.4±1.7 μM compared with the control group (A375-WT) (*P*<0.001). In addition, IC50 of M14-DR increased from 27.3±2.3 μM to 167.4±2.2 μM compared with the M14-WT (P<0.001). Subsequently, we investigated the apoptosis of the resistant strains and the control group under DTIC treatment. As shown in Fig. 1B and 1C, after 10 μg/mL DTIC treatment for 48 h, the number of apoptotic cells in A375-DR and M14-DR were both significantly reduced compared with A375-WT and M14-WT, respectively. The above experiments showed that melanoma cells resistant to DTIC have been successfully obtained in this study. Meanwhile, it also indicated that melanoma will indeed develop drug resistance under long-term chemical drug administration.

**Fig. 1.**
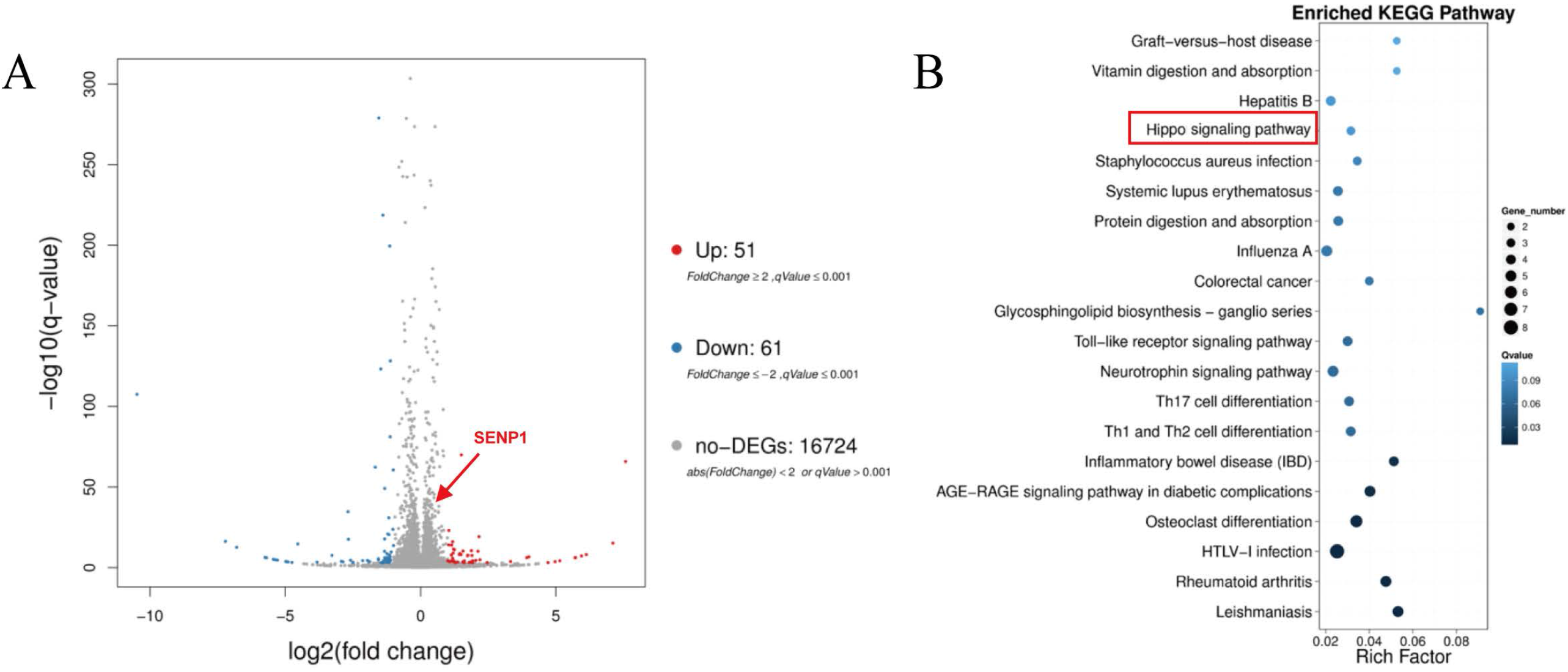
Differences between melanoma resistant cell lines and controls. (**A**) The IC50 value of drug-resistant melanoma cell lines A375-DR, M14-DR and the control group A375-WT and M14-WT against DTIC. (**B**) Anti-DTIC apoptosis ability of A375-DR, M14-DR and the control group of A375-WT and M14-WT via flow cytometry detection. (\*\*\**P*<0.001).

### Comparative transcriptome revealed differentially expressed genes in drug-resistant melanoma cells

To further explore the genes and signaling pathways involved in the tolerance mechanism in DTIC-resistant cells, we performed transcriptome analysis targeted the drug-resistant melanoma cell lines A375-DR and the according control groups A375-WT (each sample containing 3 biological replicates). As shown in **Fig. 2A**, transcriptomic analysis found that compared with the control group, there were 51 significantly up-regulated and 61 significantly down-regulated genes in melanoma drug-resistant cell lines (fold change≥1.5, *P*<0.05). Small ubiquitin-like modifier (SUMO) proteins are a family of small proteins that are covalently attached to and detached from other proteins in cells to modify their function. Among them, *SENP1* is a key protease required in the process of SUMOylation and de SUMOylation, affecting the cell cycle, cell proliferation and apoptosis status, which is also the most deeply studied SUMO proteins closely related to tumors (Zhang et al., 2021). Transcriptomics found that the *SENP1* was significantly up-regulated in the drug-resistant cell line (shown by the red arrow in **Fig. 2A**). Clustering analysis of differential genes by KEGG revealed that differential genes clustered in multiple metabolic pathways (**Fig. 2B**). Among them, the Hippo signaling pathway is involved in the regulation of key processes such as cell growth, proliferation, survival, migration and differentiation, playing an important role in organ development and the maintenance of internal environment homeostasis (Han, 2019). Among them, the key gene *YAP* of the Hippo signaling pathway was also significantly up-regulated in DTIC-resistant cell lines. This suggests that the expression change of SENP1 may be related to the abnormal activation of Hippo pathway.

**Fig. 2.**
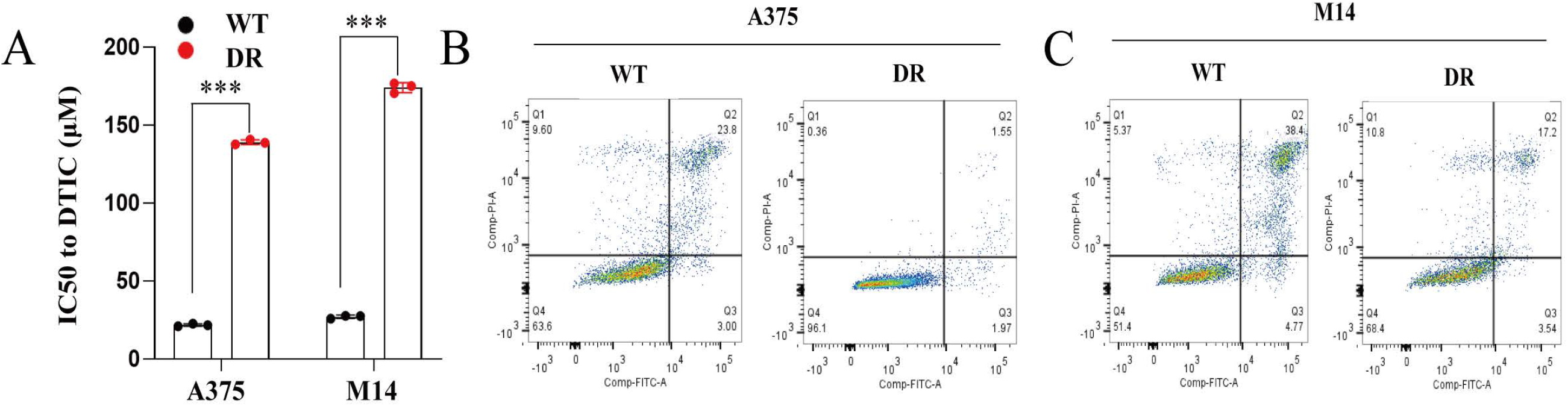
Transcriptome RNA-seq results of control and drug-resistant A375-DR cells. (A) Volcano plot analysis of differential genes, in which blue color represented the down-regulated genes while red color represented the up-regulated gene. (B) Cluster analysis of differential genes by KEGG.

### Validation of genes potentially related with drug resistance

To further verify the important genes and signaling pathways identified by transcriptomics that may be related to drug resistance of melanoma, we further used RT-qPCR and WB analysis to verify the expression of *SENP1* and *YAP* in obtained cell lines. As shown in **Fig. 3A** and **3B**, the expression levels of *SENP1* and *YAP* of the resistant strains were both significantly up-regulated in A375-DR and M14-DR, respectively. Detailly, compared with the control group A375-WT, the A375-DR had a 2.2±0.12-fold increase in the relative expression of *SENP1* (*P*<0.001) and a 0.7±0.1-fold increase in the relative expression of *YAP* (*P*<0.01). In addition, compared with the control group M14-WT, the relative expression of SENP1 in the M14-DR increased by 1.7±0.14 times (*P*<0.001) and the relative expression of YAP increased by 2.5±0.18 times (*P* <0.01). WB detection also confirmed similar conclusions (**Fig. 3C**). The above results suggested that the expression levels of SENP1 and YAP in drug-resistant cell lines are indeed increased, and may contribute to the drug resistance of melanoma.

**Fig. 3.**
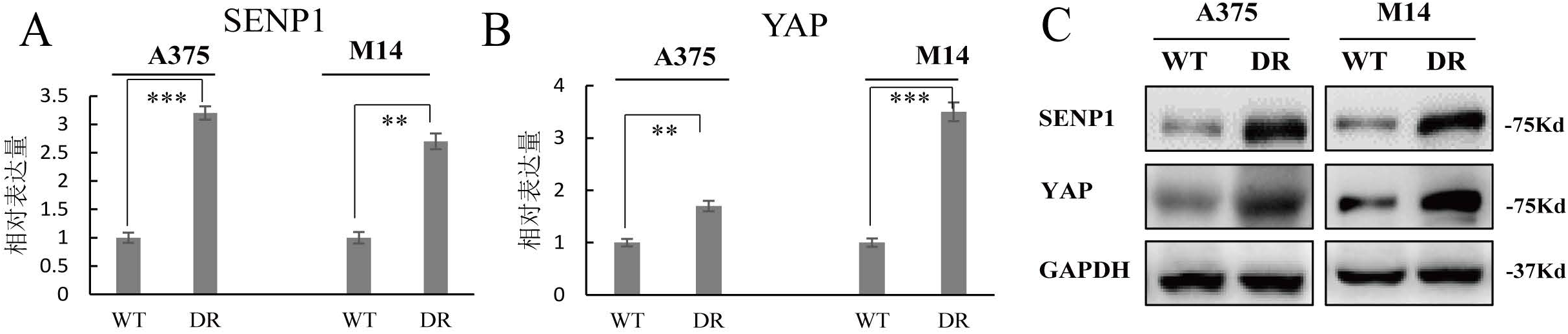
RT-qPCR and WB analysis of the expression levels of SENP1 and YAP in different cell lines. (A) The relative expression of *SENP1* in the drug-resistant melanoma cell lines A375-DR and M14-DR and the control groups A375-WT and M14-WT. (B) The relative expression of *YAP* in the drug-resistant melanoma cell lines A375-DR and M14-DR and the control groups A375-WT and M14-WT. (C) WB detection of the expression levels in the drug-resistant melanoma cell lines A375-DR and M14-DR and the control group A375-WT and M14-WT. (\*\**P*<0.01, \*\*\**P*<0.001).

### Overexpression of SENP1 in normal A375 cell improved cell viability against DTIC

Finally, to investigate the relationship between the key gene *SENP1* and drug resistance, we tried to overexpress it in normal A375 cell lines (i.e., A375-SEOE). Meanwhile, A375 containing a control plasmid was used as the blank group (i.e., A375-CT). First, WB was utilized to indicate the expression level of SENP1 in A375-CT and A375-SEOE. As shown in **Fig. 4A**, SENP1 was significantly up-regulated in A375-SEOE than that in A375-CT, demonstrating its successful overexpression. Further, viability of A375-CT and A375-SEOE toward DTIC with different concentrations was finally measured. As expected, though the viability decreased with the increasing DTIC concentration in both A375-CT and A375-SEOE, it was clearly higher in A375-SEOE than that in A375-CT especially when concentration of DTIC below 2 μM. Our results here indicated that *SENP1* indeed contributed to the drug tolerance though its detail relationship with other pathways like Hippo signal pathway remained elucidated in the future.

**Fig. 4.**
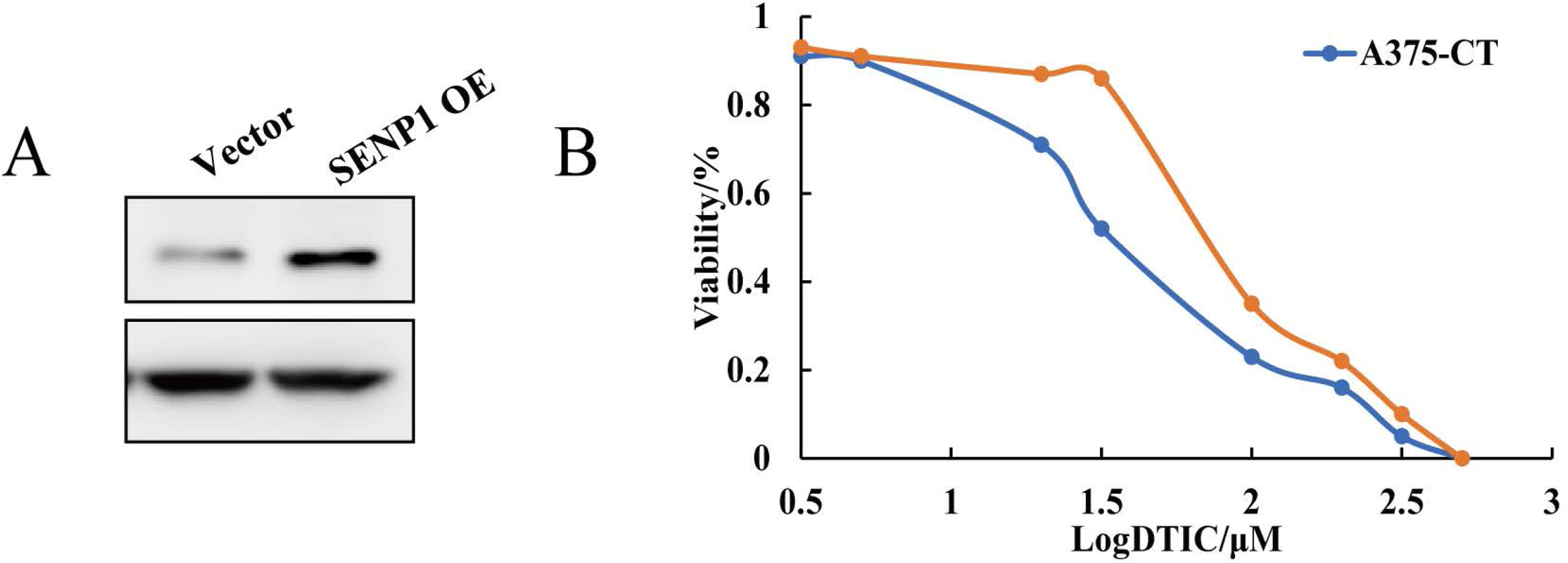
Characterization of overexpression of SENP1 in normal A375 cell. (A) WB detection of the expression levels in A375-CT an A375-SEOE. (B) Viability of A375-CT an A375-SEOE against DTIC with different concentrations.

## Discussion

Melanoma arises from genetic mutations in melanocytes and can be found in the skin, eyes, inner ear and pia mater (Gray-Schopfer et al., 2007). In addition, Melanoma accounts for approximately 1% of all skin malignancies and is the most aggressive and deadly form of skin cancer (Domingues et al., 2018). For patients with stage I-IIIB melanoma, surgery is the main treatment method along with chemotherapy, radiotherapy, immunotherapy, targeted therapy, etc. (Miller et al., 2019). Besides the adverse reactions of skin and digestive tract toxicity due to the lack of specificity of drugs (Li et al., 2015), the drug resistance of lesions to chemotherapy is also the current dilemma in the treatment of melanoma (Austin et al., 2017). Therefore, studying the causes of drug resistance in melanoma is of great significance for improving the efficacy of chemotherapy.

Protein ubiquitination modification is an important way to maintain the homeostasis of substrate proteins, which has an important regulatory effect on protein-protein interactions, subcellular localization, gene transcription activity and target protein stability (Wang et al., 2018;Jia et al., 2020). Previous studies have shown that *SENP1* is related to the tumor hypoxic microenvironment, tumorigenesis and metastasis as well as the regulation of multiple signaling pathways (Song et al., 2021;Zhu et al., 2021). In addition, studies have shown that *SENP1* is associated with tumor drug resistance. For example, Gao et al. found that SENP1 was aberrantly overexpressed in lung cancer cells, whose overexpression had a protective effect on alastine or cisplatin-treated lung cancer cells (Gao et al., 2022). In addition, Chen et al. found that, SENP1 protein accumulated in large quantities in the constructed human colon cancer cell lines resistant to irinotecan, knockdown of whom reduced the migration ability of cancer cells (Chen et al., 2021). In this study, it was found that the expression level of SENP1 was significantly increased in DTIC-resistant melanoma cells. Combined with previous reports, it is likely to play an important role in the drug resistance of melanoma.

The Hippo signaling pathway first discovered in *Drosophila* is highly conserved in mammals (Ma et al., 2019). Loss or overactivation of the Hippo pathway may lead to abnormal cell growth and tumorigenesis. In previous studies, researchers found that genes related to the Hippo pathway are often abnormally expressed in various cancers such as liver cancer, colorectal cancer, and lung cancer, and are related to the occurrence and development of tumors (Han, 2019;Xiao and Dong, 2021). The core components of the mammalian Hippo signaling pathway include cytoplasmic kinase module and a nuclear transcriptional module. Among them, YAP belonging to the nuclear transcriptional module has the function of promoting cancer (Zanconato et al., 2016). In this study, transcriptomics identified that the Hippo pathway was enriched in melanoma resistant cell lines, indicating that the signaling pathway was significantly abnormally expressed. Among them, the up-regulation of YAP was verified by RT-qPCR and WB. Combined with the abnormal expression of SENP1, whether SENP1 gene is involved in the regulation of Hippo signaling pathway, especially the regulation of YAP, is worthy of further exploration in future studies.

## Conclusions

Exploring the mechanisms of drug resistance in melanoma is of great importance for modifying the efficacy of chemotherapy in the future. In this study, via performing comparative transcriptomic between drug-resistant and parent melanoma cells, we identified the differentially expressed genes SENP1 and YAP related with Hippo signaling pathway. Notably their overexpression in normal melanoma cells also led to drug resistance. In the following work, the deep mechanisms of SENP1 and YAP participating in the drug resistance are worthy of elucidation.

## Author Contributions

BZ designed the study and analyzed the data. BZ, YL, and XT realized the experiments. BZ and SC wrote the manuscript.

## Acknowledgement

The study was supported by grants from the Natural Science Foundation of Sichuan Province (No. 2023NSFSC1549) and Sichuan Medical Association (youth innovation) Scientific Research Project (No. Q21043).

## Informed Consent Statement

There are no patents resulting from the work reported in this manuscript.

